# Speeding up eQTL scans in the BXD population using GPUs

**DOI:** 10.1101/2020.06.22.153742

**Authors:** Chelsea Trotter, Hyeonju Kim, Gregory Farage, Pjotr Prins, Robert W. Williams, Karl W. Broman, Śaunak Sen

## Abstract

The BXD recombinant inbred strains of mice are an important reference population for systems biology and genetics that have been full sequenced and deeply phenotyped. To facilitate interactive use of genotype-phenotype relations using many massive omics data sets for this and other segregating populations, we have developed new algorithms and code that enables near-real time whole genome QTL scans for up to 1 million traits. By using easily parallelizable operations including matrix multiplication, vectorized operations, and element-wise operations, we have decreased run-time to a few seconds for large transcriptome data sets. Our code is ideal for interactive web services, such as `GeneNetwork.org`. We used parallelization of different CPU threads as well as GPUs. We found that the speed advantage of GPUs is dependent on problem size and shape (number of cases, number of genotypes, number of traits). Our results provide a path for speeding up eQTL scans using linear mixed models (LMMs). Our implementation is in the Julia programming language.

## Introduction

The BXD family is an important set of approximately 150 recombinant inbred strains for which there is a large number of massive omics data sets. All members of the family have been densely genotyped and even fully sequenced. Thus any new omics data can be immediately used for quantitative expression trait locus (QTL or eQTL) mapping, and for association analyses with previously-collected phenotypes. For omic data sets collected using high-throughput technologies, additional analyses, such as transcriptional network construction or causal mediation analyses, are also practical.

The open-source GeneNetwork web service (www.genenetwork.org) (<u>1</u>) (<u>2</u>) (<u>3</u>) facilitate systems genetics and mapping by providing a searchable and exportable database of phenotypes and genotypes for a variety of organisms (including mouse, rat, and *Arabidopsis*). It also provides a suite of interactive tools for browsing the data, generating QTL maps, correlational analyses, network construction, and genome browsing. Our goal here is to develop a real-time backend for GeneNetwork to perform real-time eQTL analysis of tens-of-thousands of omics traits using key populations such as the BXDs.

To perform eQTL scans in the BXD family, one has to simply perform as many genome scans as there are phenotypes. This can be done in an “embarrassingly parallel” fashion by using standard algorithms for QTL analysis, such as those employed by R/qtl (<u>4</u>). In practice, this is too slow. For example, by using the Haley-Knott algorithms (<u>5</u>) using genotype probabilities on batches of phenotypes with the same missing data pattern, instead of using the EM algorithm (<u>6</u>) on each phenotype individually, substantial speedups are possible. This is a well-known trick and is used by R/qtl. Additionally, if only additive effects are tested, or if the population has only two genotype categories (as in a backcross or recombinant inbred line), then matrix multiplication can be used to perform Haley-Knott regression. With pre-processed data, the computation intensive part of QTL analysis can be expressed with matrix multiplication (<u>7</u>).

Processing large data set has been a challenge for genome scans. We were blessed with Moore’s Law for decades, but the central processing unit (CPU) is reaching to its physical limit of transistors. Graphical processing units (GPUs), originally used as an image processing component of a computer, have shown some compelling results to accelerate computation in various fields. General purpose graphics processor units (GPGPUs) became popular in the early 2000s because of their ability to natively handle matrix and vector operations. Such power is attractive to the scientific computing community. Zhang et al. (<u>8</u>) used GPU to simultaneously dissect various genetic effects with mixed linear model. Chapuis et al. (<u>9</u>) utilized GPU to offset heavy computation to deploy various ways for a more precise calculation of a QTL detection threshold. By using GPU-backed machine learning libraries such as TensorFlow, Taylor-Weiner et al. (<u>10</u>) reimplemented QTL mapping and Bayesian non-negative matrix factorization, achieving greater than 200 fold speedup compared to CPU versions. The ease of using such libraries has motivated the development of new methods for genomic research.

We sought to build upon these efforts aiming to perform realtime eQTL scans for the BXD populations using both CPU and GPU systems. Since programming for GPUs is often non-trivial, needing the use of low-level languages such as C++, we used the Julia programming language that offers GPU programming capabilities while retaining the simplicity of a high-level language such as R or MATLAB. Finally, since most phenotype-marker associations are null, we examined the impact of storage precision, and of only returning the highest association (LOD) score for each trait instead of a matrix of LOD scores for every pair of marker and phenotypes. We have achieved computing speeds to the extent that almost all response latency is now related to data transfer and browser display, rather that the computation. This makes real-time eQTL scans practical for the BXDs and many other well structured populations.

## Methods and Data

### A. Information about Datasets

We downloaded the BXD Genotype Database (GN Accession: GN600) and two sets of transcriptome data, UTHSC Affy MoGene 1.0 ST Spleen (GN Accession: GN283) and UMUT Affy Hippocampus Exon (GN Accession: GN206) from GeneNetwork to build and refine methods. The genotype file includes 7321 markers by 198 BXD strains; the spleen dataset has data for 79 BXD strains and for 35556 transcripts, while the hippocampus dataset has data for 70 BXD strains and 1,236,087 probe sets. Data cleaning and wrangling was performed using R/qtl (<u>4</u>) and R/qtl2 (<u>11</u>).

### B. Linear Model

Let *y*_*i*_ denote a vector for the *i*-th expression trait for *n* individuals in the (*i* = 1, …, *m*). We define a univariate linear model as follows:

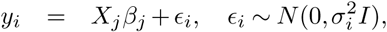

where *X*_*j*_ is a matrix including the *y*-intercept and the *j*-th candidate genetic marker (*j* = 1, …, *p*) without any co-variate(s), *β*_*j*_ is a vector of the *j*-th eQTL effects, and *ϵ*_*i*_ is random error distributed as 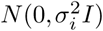. Suppose *RSS*_0*i*_ is the residual sum of squares under the null hypothesis of no eQTL, and *RSS*_1*ij*_ is the residual sum of squares under the alternative of existing eQTL at the *i*-th trait and *j*-th genetic marker. Then, the *LOD*_*ij*_ score for a one-df test can be written as:

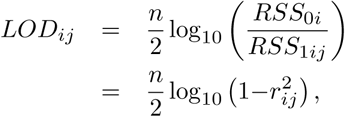

where *r*_*ij*_ is the correlation between the *i*-th expression trait and *j*-th marker. If *Y* ^∗^ and *G*^∗^ are respectively standardized trait (*Y*) and genotype (*G*) matrices, then the correlation matrix is simply

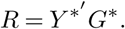

Since matrix multiplication is a parallelizable operation for which optimized routines are available, this formula is very atrractive for bulk calculation of LOD scores. The formula can be extended for LOD scores adjusted by covariates. The idea is to project genetic markers and gene expressions onto the space orthogonal to the covariates and to compute the corresponding correlation matrix just as we did for the no-covariate case. In other words, let *Z* be a matrix of covariates including *y*-intercept. The projection orthogonal to the co-variate space is then *P* = *I* −*Z*(*Z′Z*)^−1^*Z′*. The genotype matrix (*G*) and gene expressions (*Y*) are now transformed into *G*_*z*_ = *PG, Y*_*z*_ = *PY*, respectively. This is the same as calculating the residuals after regressing on *Z*. Standardizing them 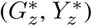 followed by multiplication yields the correlation matrix (*R*_*z*_) just as shown above. Figure 1 gives a visual representation of the matrix multiplication.

**Fig. 1.**
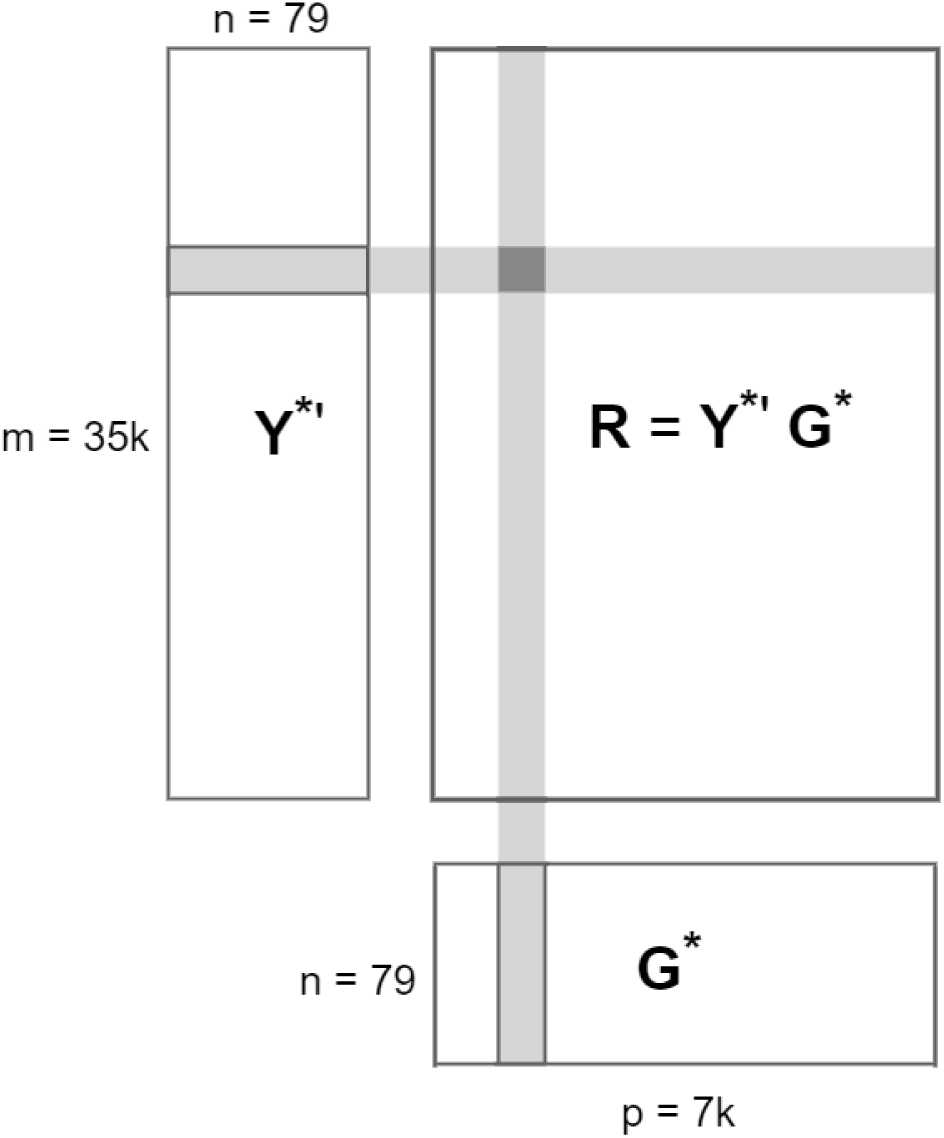
Schematic of data and correlation calculation: *Y* is an expression phenotype matrix, *G* is a genotype matrix, and *R* = *Y* ^∗′^*G*^∗^ is a correlation matrix. LOD scores are a function of the correlation matrix.

### C. Acceleration Techniques

While it is true that many programs can achieve tens or even hundreds of speedup by utilizing GPUs, the difference needs to be examined with more care. Sometimes, the reported CPU time is using single thread, and multithreaded CPU time may bring the performance gap of CPU and GPU closer than claimed. Also depending on the library chosen for CPU, speed might vary depending on whether the library is optimized for such computation or hardware. We believe for a fair comparison, both CPU and GPU functions should be optimized at maximum performance and count in all necessary overhead. The following section explains our efforts of optimization on CPU and GPU functions.

#### C.1. Multithreaded CPU operations

Our goal is to build a backend for GeneNetwork that allows researchers to interact with data in real time. Such requirement sets up the standard that the genome scan must finish within seconds. To bring out the best performance of CPU, we use multithreaded operations whenever possible. Julia (<u>12</u>) (<u>13</u>), our choice of the programming language, provides simple yet safe syntax for multithreading. It is simply done by adding Threads.@thread macro to indicate to Julia that the following `for` loop is the multithreaded region. Using `Threads.nthreads()` function can show the number of threads in julia, and the default number of threads we use is 16 threads.

#### C.2. Matrix and Vectorized Operations

Since our algorithm largely depends on matrix operations, it is natural to find the fastest way to achieve the best result, regardless of computing platforms. There are various matrix libraries available for CPU, such as gslBLAS and OpenBLAS (<u>14</u>). They target different hardware or use various techniques to get optimal results. Multithreaded matrix multiplication is the default in OpenBLAS, so as not to require an extra coding effort to make the CPU version of matrix multiplication run in parallel. We therefore chose OpenBLAS as the CPU computing library.

Matrix multiplication and element-wise operations are algorithmically free of data and function dependency, so that they are amendable to GPU’s parallel computing power. Julia provides various packages for GPU including CUDA (<u>15</u>) bindings. Our chosen hardware for GPU is from Nvidia, which requires its proprietary library, CUDA. Besides the hardware requirement, CUDA is also mature and has been recognized in the scientific computing community. https://www.overleaf.com/project/5eea5c1d4f553e00016a0611 *cuBLAS* (<u>16</u>) library provides a fast GPU implementation of the BLAS from Nvidia. For matrix operations on GPU, we used the cuBLAS library.

Not only do we want to see how much speed up we can get from using GPU, but we also experiment whether the shape of matrix will affect the speedup. Of course, most of the time, one cannot pick the size and shape of data in a matrix form, but such information would help researchers as a rough guidance of whether it is worth considering the GPU option before investing programming efforts for GPU. We ran matrix multiplication with different shapes of matrices and compared the running time of CPU and GPU. CPU time is measured by matrix multiplication from the OpenBLAS library using 16 threads. GPU time includes all overhead of using GPU, which involves device launch, data transfer and all necessary API calls. In order to make a fair comparison between CPU and GPU, we need to use maximum strength of both and include all necessary cost.

The experiment setup is to multiply two input matrices, A(m *×* n), and B(n *×* p), and produce an output matrix C (m *×* p). The range of *m, n* and *p* is between 2^4^ and 2^17^ in log scale. Due to the GPU hardware limitations, we can only compare the result when the size of input and output (I/O size) matrices, in total, is less than 16 GB. The result is shown in Figure 2. The *x*-axis of Figure 2 is the dimensions of matrix whose I/O size is over 12 GB. This is because using GPU involves a lot of overhead. Such overhead can only be justified by processing large data sets. The *y*-axis is the speedup of GPU compared with CPU in log scale. From this figure, matrices whose shapes are closer to square matrices get better speedups from GPU. Matrix multiplication is up to 5 times faster on GPU than that on 16 threaded CPU.

**Fig. 2.**
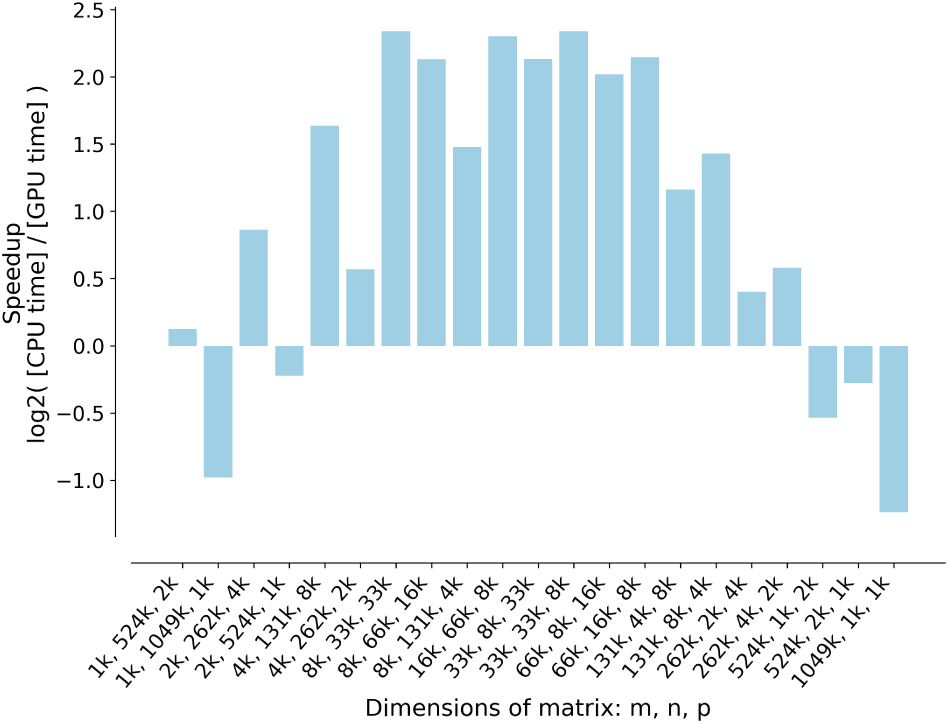
Variation of GPU vs CPU speedup with matrix shape.

#### C.3. Single precision

Precision means the smallest difference between two representable numbers. Floating point numbers, in scientific computing, are usually stored in double precision. Double precision floating point numbers take up 8 bytes in memory while single precision takes up 4 bytes. In addition to the difference in storage size, the speed for calculation using single and double precision also varies by hardware. For example, the GPU throughput (the number of floating point calculation per second, measured in FLOPS) for double precision is 1/32 of single precision on a Nvidia GTX 1050 GPU, and 1/4 on a Nvidia Tesla K80. Differences in optimization techniques, underlying architecture cause such performance gap. Considering the storage size, data transfer speed and throughput, single precision brings multiple benefits when precision is not the priority concern.

#### C.4. GPU operations

Originally used in graphics, GPU has taken off as a general computing device in recent years because of its massive number of cores at a lower price range, and fast GPGPU libraries such as CUDA and OpenCL. Based on our profiling result, the time consuming parts of our genome scan method are matrix multiplication and element-wise operations. Both are amenable to GPU heterogeneous computing architecture since they have no data race conditions and low data dependencies. However, GPU also has its own limitations. To truly utilize the maximum computing power of GPU, one needs to think creatively to work around such limitations from GPU. For example, during our experiment, memory transfer between host and device is really slow. Profiling the result shows that 98% of total genome scan time is spent on memory transfer. To cope with this limitation, instead of offloading the entire correlation matrix, we use GPU to calculate the max LOD score of each expression trait and output the maximum.

#### C.5. Julia language

Although a programming language cannot really be classified as an optimization technique, the choice of programming language can affect the speed of the program. Ease of use is also considered because the cost of any program comprises not only recurring expense of running time on a piece of hardware but also the front load expense of programming effort. Shorter development time is valuable because researchers can quickly prototype and experiment with different models, but performance should also be equally important, if not more. Our language choice is Julia.

## Results

Our method uses the linear model and simplified the genome scan process to basic matrix operations. The timing of our method is shown in Table 1. The execution time without finding the maximum LOD score of phenotype (full results) using double precision is 0.55 second if we only use CPU. Getting the same result using the same dataset from R/qtl took about 36.5 seconds.

**Table 1.**
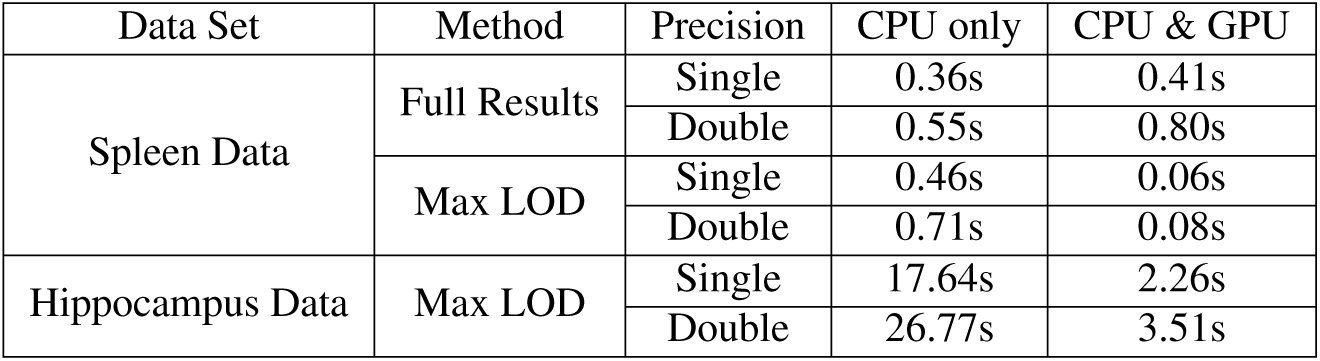
Time comparison between CPU and GPU.

### D. Benefit of using single precision

Table 1 also shows the execution time using single and double precision. In all cases, genome scan runs faster using single precision than using double precision. Using single precision provides benefits in 3 aspects: memory storage, data transfer, and arithmetic calculation.

### E. Benefit of using GPU

In parallel computing, Amdahl’s law is used to find the theoretical maximum speedup by improving a particular part of a program. For example, if a program takes ten minutes for a serial processor, and a function that takes nine hour of those ten minutes can be parallelized, and then, the theoretical speedup, no matter how many processors are used, can not be more than ten times because the minimum execution time of this program is one minute. Therefore, profiling the entire genome scan process is a prerequisite for the optimization. Often, profiling would consider space and time complexity. Our primary concern is the time taken by each function, and therefore only timing information is considered in our profiling. We used Julia’s built-in Sampling Profiler to find our target function for GPU because it is less intrusive than the other profiling methods. The genome scan process includes the following steps:

- Calculate standardized matrices (*G*^∗^, *Y* ^∗^) for input matrices (*G, Y*)
- Get a correlation matrix (*R*) by multiplying the standardized matrices
- Calculate LOD scores

Our profiling result shows that the second and third steps take up 90% of the time and involve parallelizable matrix operations. Hence, they are our candidates for GPU acceleration. To make a fair comparison of CPU and GPU, the timing shown in Table 1 is the total execution time for genome scan. There is overhead of using GPU, such as data transfer and setting up context. The timing shown for CPU & GPU here included all overhead for fairness. We ran the genome scan process 10 times and chose the median to remove the randomness of each run and warm-up time of GPU. The *Full Results LOD* method shows the timing when running the scan with *CPU only* and *CPU&GPU*. As the table shows, the CPU&GPU combination did not show any improvement over the serial version (CPU only) in terms of the performance time. That is, the former is rather slower than the latter regardless of precision for the data type. We used a GPU profiler *nvprof* (<u>17</u>) to investigate what is the bottleneck of GPU. The result is, 98% of the time is spent on data transfer from GPU to CPU (device to host). As shown in Figure 1, the input matrices Y’ and G are small compared with the output matrix R. For the BXD spleen dataset, Y’ matrix is 17 MB, G matrix is 21 MB, but R matrix is about 4GB. Data offloading is the main bottleneck for our GPU implementation. To cope with this limitation of GPU, we further developed *Max LOD* method because the main interest of genome scan is to find the maximum LOD score of every phenotype. This step is highly parallelizable, can utilize GPU’s massive cores, and, at the same time, reduces the amount of data that needs to be transferred back to host. Table 1 shows that *Max LOD* method for the spleen data reduces CPU&GPU execution time down to 0.079 seconds when using double precision. Compared with CPU only, which took 0.55 seconds, this GPU modification provides 7 times speedup. As we mentioned earlier, since we are porting the computation to GPU, which took 90% of the total execution time, according to Amdahl’s Law, the theoretical maximum speed up is 10 times. We got 7 times speed up for genome scan using GPU. For the hippocampus data, which contains over 1 million phenotypes, the GPU *Max LOD* implementation took 3.51 seconds for double precision and 2.26 seconds for single precision. As a backend service to a website, this wait time is much more preferable than the CPU timing for real time interactions.

### F. Benefit of using Julia

In our experiment of matrix multiplication, Julia’s speed is comparable to C/C++. However, the low learning curve, clean syntax, as well as support to GPU programming libraries such as CUDAnative (<u>18</u>) and CuArrays (<u>19</u>) affords much lower programming efforts than C/C++. Compared with writing GPU functions in C/C++, writing in Julia is cleaner and easier because it requires much less boilerplate code. Below is some code snippets to demonstrate GPU code, calling a library or writing a custom kernel in Julia. The first example illustrates how to call CUBLAS from Julia, and the second example shows how to write a custom kernel in Julia. For the reason of page limitation, we will not show the corresponding C code. An example of using CuBLAS with C can be found online (<u>20</u>).

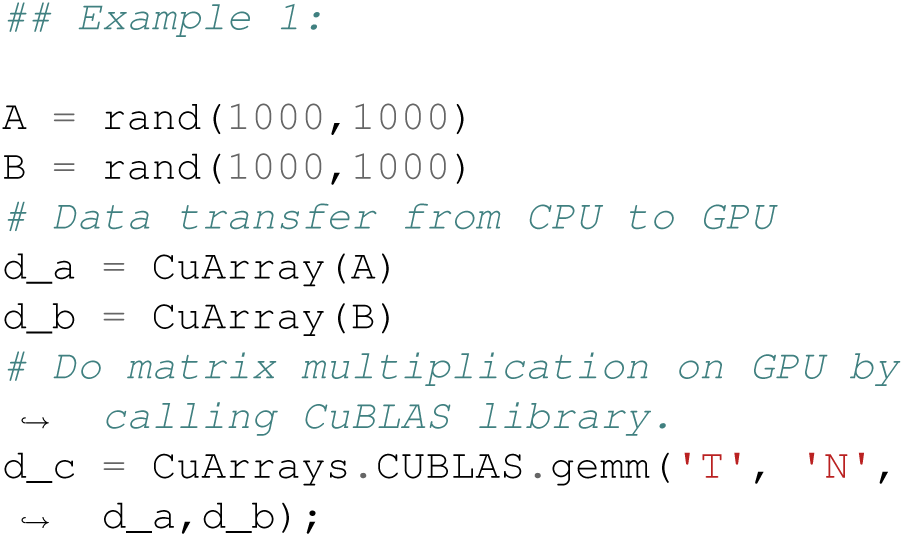

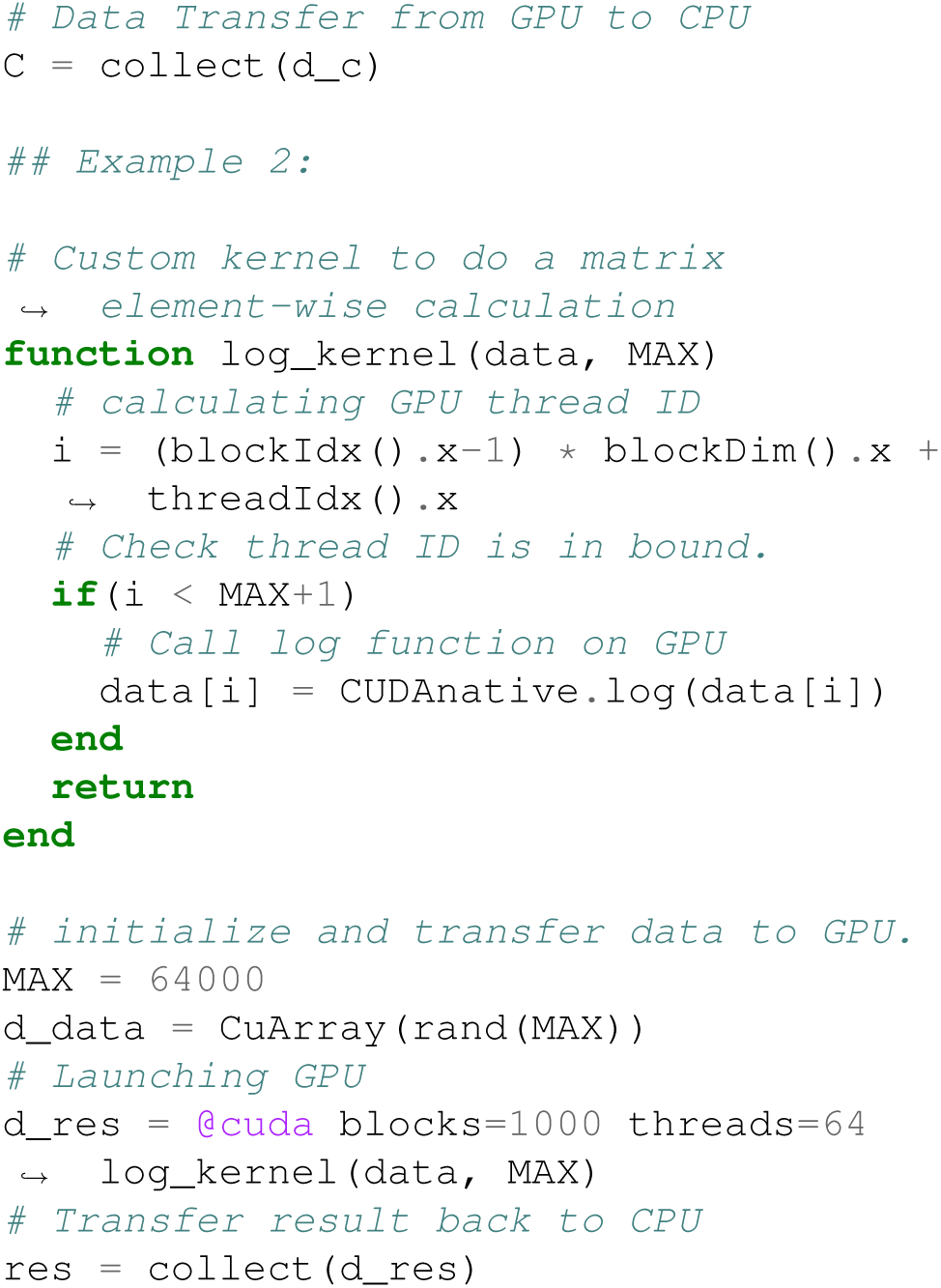

### G. Software Availability

The source code for this package is publicly available on Github as the LMGPU Julia package: https://github.com/senresearch/LMGPU.jl. This repository contains three folders: *data, src* and *test*. Using Julia language and running `test.jl` in *test* folder will run our method using either the spleen data or hippocampus data in *data* folder. Thes package uses openBLAS and CUDA10.1. To use this package, one needs to have a Nvidia GPU in the test machine, and have the Julia executable, openBLAS and CUDA libraries installed.

### H. Platform

Our platform for computation:

Hardware:

- CPU: Intel Xeon Gold 6148; 40 cores @ 2.40GHz, 192GB
- GPU: Tesla V100-PCIE; 5120 Cores @ 1380 MHz, 16GB

Software:

- Programming environment: Julia; CentOS 7
- Libraries: CUDA v10.1 and cuBLAS; OpenBLAS
- Profilers: Julia Profiler; nvprof

## Conclusion

We examined the effectiveness of using GPUs for speeding up eQTL scans in the BXD family of recombinant inbred lines. We investigated the effect of different matrix size and shape on GPU speed up for matrix multiplication. Closer to a square matrix [WHICH MATRIX IS SQUARE] wins the best speedup over CPU when matrix multiplication is implemented on GPU. With the maximum total input data size of 16 GB, matrix multiplication is up to 5 times faster on GPU than on 16 threaded CPUs. Our LMGPU package takes advantage of GPU’s massive parallel computing power, Julia’s elegant syntax, fast performance for quick prototyping, and performance-minded implementation in one language. LMGPU also uses various techniques such as multi-threaded operations and finding the maximum LOD score on GPU to achieve close to real-time genome scan. We are able to run genome scan for the spleen data in 0.06 seconds and one for the hippocampus data with over one million traits in 3 seconds. As a backend for the GeneNetwork web service, LMGPU will enable researchers to do real-time interaction with the data. It can also be used as a stand-alone Julia package for running eQTL scans.

The LMGPU package also has some limitations. Currently the best speed up of GPU uses an algorithm that finds the maximum LOD score to minimize data transfer between GPU and CPU, which was the bottleneck for our LM algorithm. For those who are interested in the whole matrix of LOD scores using the CPU alone might be the best option. LMGPU also only supports 1-degree freedom test currently, with no missing data in input dataset; any missing data has to be handled in pre-processing and will add to the computation time. Currently the the LOD scores are fit using a linear model; for many problems a linear mixed model (LMM) (<u>21</u>) is of interest. We expect to build on the current work to tackle that problem in the future.

## Bibliography

1. Zachary Sloan, Danny Arends, Karl W Broman, Arthur Centeno, Nicholas Furlotte, Harm Nijveen, Lei Yan, Xiang Zhou, Robert W Williams, and Pjotr Prins. GeneNetwork: framework for web-based genetics. The Journal of Open Source Software, 1(2), 2016.

2. Megan K Mulligan, Khyobeni Mozhui, Pjotr Prins, and Robert W Williams. GeneNetwork: a toolbox for systems genetics. In Systems Genetics, pages 75–120. Springer, 2017.

3. Elissa J Chesler, Lu Lu, Jintao Wang, Robert W Williams, and Kenneth F Manly. WebQTL: rapid exploratory analysis of gene expression and genetic networks for brain and behavior. Nature neuroscience, 7(5): 485–486, 2004.

4. Karl W Broman, Hao Wu, śaunak Sen, and Gary A Churchill. R/qtl: QTL mapping in experimental crosses. Bioinformatics, 19(7): 889–890, 2003.

5. Chris S Haley and Sarah A Knott. A simple regression method for mapping quantitative trait loci in line crosses using flanking markers. Heredity, 69(4): 315, 1992.

6. Eric S Lander and David Botstein. Mapping mendelian factors underlying quantitative traits using RFLP linkage maps. Genetics, 121(1): 185–199, 1989.

7. Andrey A Shabalin. Matrix eQTL: ultra fast eqtl analysis via large matrix operations. Bioinformatics, 28(10): 1353–1358, 2012.

8. Fu-Tao Zhang, Zhi-Hong Zhu, Xiao-Ran Tong, Zhi-Xiang Zhu, Ting Qi, and Jun Zhu. Mixed linear model approaches of association mapping for complex traits based on omics variants. Scientific reports, 5:10298, 2015.

9. Guillaume Chapuis, Olivier Filangi, Jean-Michel Elsen, Dominique Lavenier, and Pascale Le Roy. Graphics processing unit–accelerated quantitative trait loci detection. Journal of Computational Biology, 20(9): 672–686, 2013.

10. Amaro Taylor-Weiner, Francois Aguet, Nicholas Haradhvala, Sager Gosai, Shankara Anand, Jaegil Kim, Kristin Ardlie, Eliezer Van Allen, and Gad Getz. Scaling computational genomics to millions of individuals with GPUs. bioRxiv, page 470138, 2019.

11. Karl W Broman, Daniel M Gatti, Petr Simecek, Nicholas A Furlotte, Pjotr Prins, śaunak Sen, Brian S Yandell, and Gary A Churchill. R/qtl2: software for mapping quantitative trait loci with high-dimensional data and multiparent populations. Genetics, 211(2): 495–502, 2019.

12. Jeff Bezanson, Alan Edelman, Stefan Karpinski, and Viral B Shah. Julia: A fresh approach to numerical computing. SIAM review, 59(1): 65–98, 2017.

13. Julia 1.0. https://julialang-s3.julialang.org/bin/linux/x64/1.0/julia-1.0.5-linux-x86_64.tar.gz, 2018. Accessed: 2020-06-19.

14. Qian Wang, Xianyi Zhang, Yunquan Zhang, and Qing Yi. AUGEM: automatically generate high performance dense linear algebra kernels on x86 CPUs. In SC’13: Proceedings of the International Conference on High Performance Computing, Networking, Storage and Analysis, pages 1–12. IEEE, 2013.

15. John Nickolls, Ian Buck, Michael Garland, and Kevin Skadron. Scalable parallel programming with CUDA. Queue, 6(2): 40–53, March 2008. ISSN 1542-7730. doi: 10.1145/1365490.1365500.

16. NVIDIA. cuBLAS. https://developer.nvidia.com/cublas, 2019. Accessed: 2019-07-24.

17. NVIDIA. nvprof. https://docs.nvidia.com/cuda/profiler-users-guide/index.html, 2019. Accessed: 2019-07-24.

18. Tim Besard, Christophe Foket, and Bjorn De Sutter. Effective extensible programming: Unleashing Julia on GPUs. IEEE Transactions on Parallel and Distributed Systems, 2018. ISSN 1045-9219. doi: 10.1109/TPDS.2018.2872064.

19. Tim Besard. cuArray. https://github.com/JuliaGPU/CuArrays.jl, 2019. Accessed: 2019-07-24.

20. Nvidia. CuBLAS example code. https://docs.nvidia.com/cuda/cublas/index.html#example-code, 2019. Accessed: 2019-10-28.

21. Christoph Lippert, Jennifer Listgarten, Ying Liu, Carl M Kadie, Robert I Davidson, and David Heckerman. FaST linear mixed models for genome-wide association studies. Nature methods, 8(10): 833, 2011.

